# *Gelsolin* and *dCryAB* act downstream of muscle identity genes and contribute to preventing muscle splitting and branching in *Drosophila*

**DOI:** 10.1101/784546

**Authors:** Benjamin Bertin, Yoan Renaud, Teresa Jagla, Guillaume Lavergne, Cristiana Dondi, Jean-Philippe Da Ponte, Guillaume Junion, Krzysztof Jagla

## Abstract

A combinatorial code of identity transcription factors (iTFs) specifies the diversity of muscle types in *Drosophila*. We previously showed that two iTFs, Lms and Ap, play critical role in the identity of a subset of larval body wall muscles, the lateral transverse (LT) muscles. Intriguingly, a small portion of *ap* and *lms* mutants displays an increased number of LT muscles, a phenotype that recalls pathological split muscle fibers in human. However, genes acting downstream of Ap and Lms to prevent these aberrant muscle feature are not known. Here, we applied a cell type specific translational profiling (TRAP) to identify gene expression signatures underlying identity of muscle subsets including the LT muscles. We found that *Gelsolin* (*Gel*) and *dCryAB*, both encoding actin-interacting proteins, displayed LT muscle prevailing expression positively regulated by, the LT iTFs. Loss of *dCryAB* function resulted in LTs with irregular shape and occasional branched ends also observed in *ap* and *lms* mutant contexts. In contrast, enlarged and then split LTs with a greater number of myonuclei formed in *Gel* mutants while *Gel* gain of function resulted in unfused myoblasts, collectively indicating that *Gel* regulates LTs size and prevents splitting by limiting myoblast fusion. Thus, *dCryAB* and *Gel* act downstream of Lms and Ap and contribute to preventing LT muscle branching and splitting. Our findings offer first clues to still unknown mechanisms of pathological muscle splitting commonly detected in human dystrophic muscles and causing muscle weakness.

## Introduction

Diversification of cell types is a fundamental process during the development of multicellular organisms and is essential in building functional organs. The muscle network in *Drosophila* embryos, composed of 30 muscle fibres per abdominal hemisegment, offers a tractable system for studying cell diversification. Despite common characteristics such as formation by myoblast fusion and the capacity to contract, each embryonic *Drosophila* muscle has a specific size, orientation, number of nuclei, attachment and innervation [1]. However, how these features are acquired at the muscle-specific level remains unclear, although it is generally accepted [2, 3] that the muscle founder cells (FCs), which are at the origin of muscle fibres, harbour all the information required for individual muscle identity. This information is thought to be provided by the combinatorial expression of identity genes encoding identity transcription factors (iTFs) [4, 5]. They are all activated in subsets of muscle progenitors and/or FCs and share one important feature: their loss of function leads to the loss or aberrant properties of a muscle in which they are expressed [1]. Some identity genes such as *lateral muscles scarcer* (*lms*) [6] show a restricted expression in lateral muscles only, whereas others, such as *slouch* (*slou*) display a broader expression pattern in ventral, lateral and dorsal domains [7, 8]. Most *Drosophila* iTFs have their vertebrate counterparts, some of which (e.g. Org-1/TBX, Caup/IRX, Tup/ISL), play conserved roles during musculature specification in vertebrates [9–11]. Knowledge gained in *Drosophila* on iTFs and their downstream targets could thus cast light on how muscle diversification processes are regulated in general but also how aberrant muscle features such as branching and splitting are acquired. Whole genome approaches based on ChIP experiments have shown that iTFs directly regulate not only other identity genes, but also downstream regulators of muscle identity, the “realisator genes” [12–14]. We previously demonstrated that Eve, Lb and Slou iTFs regulate the number of fusion events by setting expression levels of genes that act as identity realisators in a muscle-specific manner [13, 15]. However, the identification of “realisator genes” has so far been limited to only a few examples, owing to the technical challenges of detecting gene expression in specific muscle populations during embryogenesis.

To further analyse diversification processes and identify genes acting downstream of iTFs, we optimised translating ribosome affinity purification (TRAP) [16] to small subsets of FCs and developing muscle precursors [17]. TRAP-purified mRNA profiling followed by bioinformatic analysis and generation of temporal transition profiles identified muscle subsetspecific translatome signatures with *Gelsolin* (*Gel*) and *dCryAB* as new identity realisator genes controlling shape- and size-related properties of Lms-expressing muscles.

Intriguingly, *dCryAB* and *Gel* act downstream of Ap and Lms and their loss-of-function phenotypes recall dystrophic muscle branching/splitting in humans [18].

## Results

### Translational profiling (TRAP) of muscle subsets identifies *dCryAB* and *Gelsolin* expressed predominantly in LT muscles

TRAP is based on the polysome capture of the GFP-tagged ribosomes with their associated mRNAs (Fig. 1A). Here we tagged polysomes with UAS-Rpl10A-GFP in two muscle subsets using Slou-GAL4 or Lms-GAL4 drivers (Fig. 1B) and in all embryonic muscles using Duf-GAL4. For each muscle subset, translational profiling was performed on embryos collected from three developmental time windows T1: 7–10 h AEL, T2: 10–13 h AEL and T3: 13–16 h AEL covering the main muscle development steps. To assess the specificity of TRAP-based muscle targeting, we analysed GO terms and found that the up-regulated genes fitted muscle-related GOs while GO categories associated with the list of down-regulated genes were not related to muscle developmental processes (Fig. 1C). Gene expression profiling with total embryonic RNA (input fractions) as a reference was used to identify differential gene expression and perform spatial and temporal gene clustering. A significant portion of upregulated genes (FC > 2, *p* < 0.05) turned out to be common to the restricted muscle populations (Slou- and Lms-positive) and the overall (Duf-positive) population (Fig. 1D). The percentage of Slou-specific transcripts remained constant at different time points, while the proportion of Lms-specific transcripts increased (Fig. 1D). This difference may be due to Slou-expressing muscles being more heterogeneous than Lms-positive ones. Finally, to test muscle type-specific gene expression, we generated volcano plots (Fig. S1A,B) and observed that genes with previously characterised expression patterns in muscle subsets displayed expected up- or down-regulation. For example, in Lms-positive muscles, *ap* and *lms* gene transcripts were enriched, whereas *slou* and *org1* transcripts, specific for Slou-positive muscles, were depleted (Fig. S1B). We also tested whether TRAP would detect “low expression” genes. To do so, we crossed our lists of enriched and depleted transcripts with modEncode datasets [19] and found that more than 30% of up-regulated TRAP-ed genes entered the “low expression” modEncode category, whereas most depleted transcripts fitted the “high expression” modEncode category (Fig. S1C,D). TRAP-based translational profiling of muscle subsets was thus specific for the targeted muscle populations and sensitive enough to detect low transcript levels.

**Figure 1.**
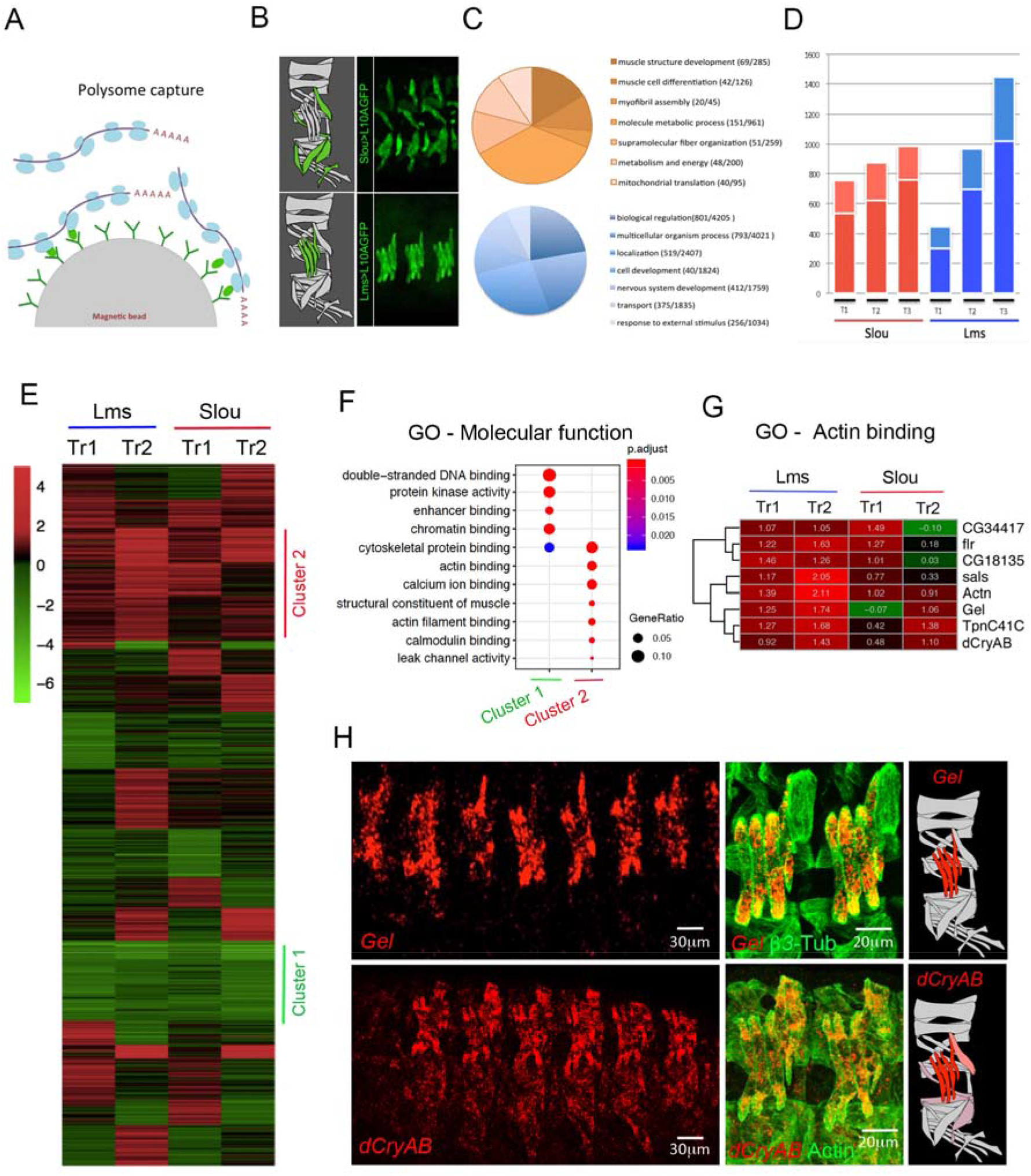
Translational profiling of embryonic muscles using TRAP. **A.** Schematic representation of cell-specific polysome capture during TRAP. **B**. TRAP-targeted muscle populations revealed by anti-GFP immunostaining of stage 16 *Slou>RpL10a-eGFP* and *Lms>RpL10a-eGFP* embryos. **C**. Pie chart showing distribution of GO terms (biological processes) for genes commonly up-regulated (orange) or down-regulated (blue) in Slou- and-Lms positive muscles and in all time windows. **D**. Bar plots showing proportions of genes whose expression is commonly up (FC>2) in individual muscle subsets and in all muscles (dark red and dark blue) versus genes up-regulated exclusively in Slou (light red) and Lms (light blue) populations. **E**. Heatmap showing the transition profiles of gene expression in Lms and Slou populations. Tr1 represents gene expression transition from the time window T1 to T2 and Tr2 the transition from T2 to T3. Cluster 1 (green) and cluster 2 (red) genes are indicated. **F**. GO comparison of genes belonging to Cluster 1 and Cluster 2. **G**. A zoomed view of the transition profile heatmap restricted to genes from cluster 2 that belong to the “actin binding” GO category. Respective Tr1 and Tr2 fold changes are indicated on the heatmap. **H**. *In situ* hybridisation showing that *Gel* and *dCryAB* that are part of the “actin binding” GO class are predominantly expressed in LT muscles. Lateral views of stage 15 embryos are shown. Anti-actin or anti-□3 tubulin antibodies are used to reveal muscle pattern. Schemes of muscles in an abdominal segment are shown with *Gel* and *dCryAB* expression indicated by a color code: red – high expression and light pink – low expression levels.

We then applied temporal transition profiling to identify clusters of genes showing similar dynamics of expression patterns, thus potentially under common upstream regulatory cues. We considered that this approach could be applied to TRAP datasets to identify novel muscle identity realisator genes acting downstream of iTFs [15]. The generated temporal transition heatmap for Slou- and Lms-positive muscles revealed clusters of genes with several expression behaviours (Fig. 1E).

Here we focused on gene clusters showing two contrasting transition behaviours, “down-down” (Cluster 1) and “up-up” (Cluster 2). Cluster 1 genes were characterised by enrichment of GOs associated with “chromatin binding” (Fig. 1F). Among Cluster 1 genes, we found *twist* and several Notch pathway-involved genes whose expression needs to be turned down while muscle differentiation progresses. By contrast, Cluster 2 showed a significant enrichment of several GO terms related to muscle development (Fig. 1F) including conserved genes encoding actin-binding proteins: *CG34417/Smoothelin, flr/WDR1, CG18135/Gpcpd1, sals/SCAF1, Actn/ACTN2, Gel/GSN, TpnC41C/CALM1, dCryAB/l(2)efl/CRYAB* (Fig. 1G). We found this sub-cluster of particular interest as it contains genes with similar biological functions and differential Lms-versus Slou-transition profiles. Among them, *Gel* and *dCryAB* with “up-up” transition profile in the Lms subpopulation (Fig. 1G) are both preferentially expressed in Lms-positive LT muscles (Fig. 1H). *Drosophila Gel* belongs to the conserved Gelsolin/Villin family of actin interactors [20] with actin depolymerisation activity [21, 22], whereas *dCryAB* codes for a small heat shock protein (sHSP) carrying an actin-binding domain and known to interact with cheerio/filamin [23]. *Gel* transcripts could be detected in LT muscle precursors from late stage 14 (Fig. 1H) but not earlier (Fig. S2A,A’,B,B’). Moreover, in late stage embryos *Gel* is prominently expressed in the visceral muscles and developing fat body (Fig. S2C,C’). *dCryAB* transcripts accumulate preferentially in LTs and in DT1 starting from early embryonic stage 15 (Fig. 1H) and at a lower levels in a larger population of lateral and ventral muscles including SBM and VT1 (Fig. 1H, Fig. 2A,A’). LT muscle specification is under the control of identity gene *Msh* and downstream iTFs, Ap and Lms [6]. *Gel* and *dCryAB* LT expression (Fig. 2A-B) is dramatically reduced in *lms* (Fig. 2CD) and in particular in *lms/ap* mutant embryos (Fig. 2E-F) and *Gel* ectopically activated by panmuscular Ap (Fig. 2G,H) showing that LT iTFs positively regulate *Gel* and *dCryAB* (Fig. 2I).

**Figure 2.**
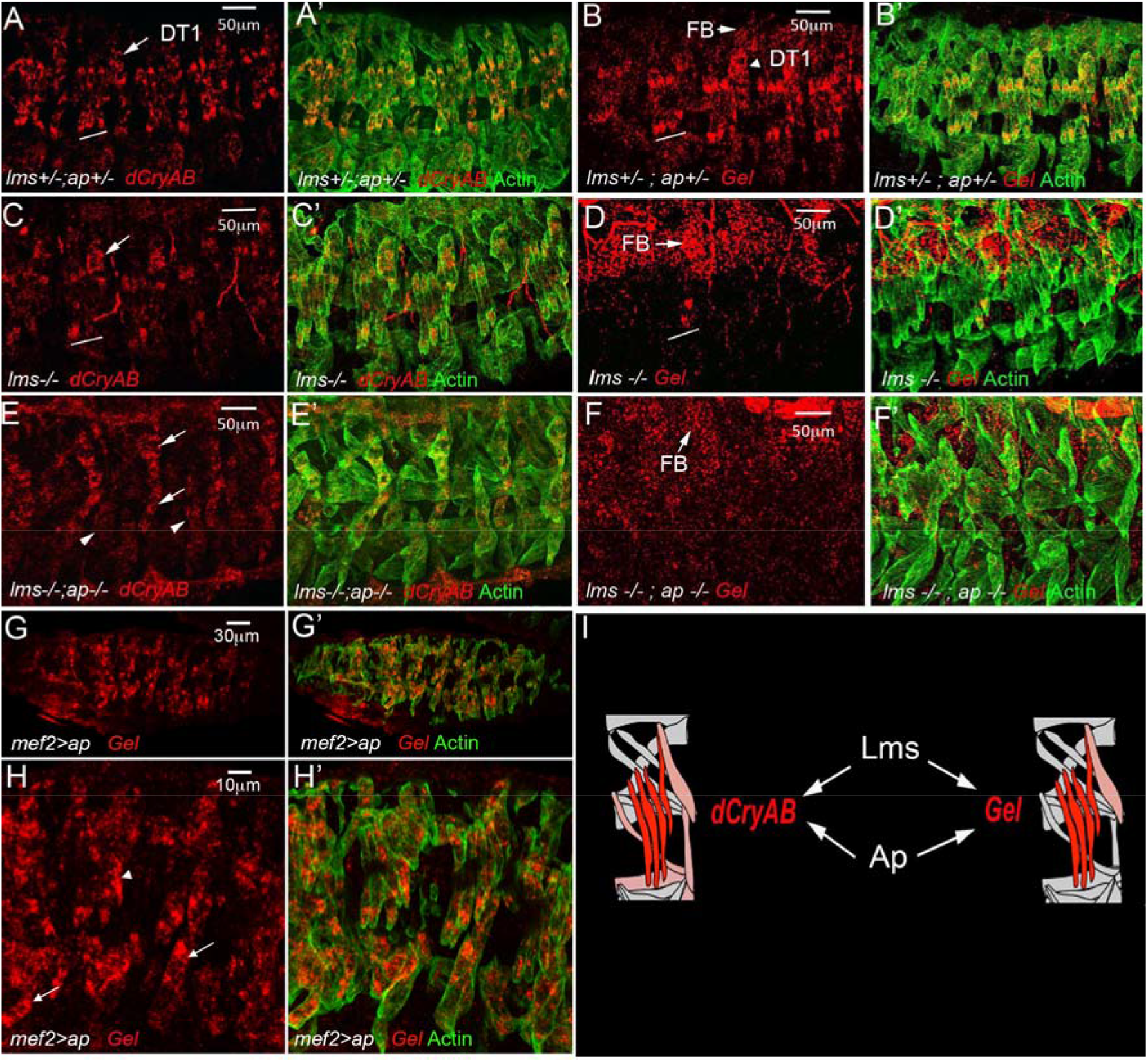
Lms and Ap positively regulate the expression of *dCryAB* and *Gel* in LT muscles.’ *dCryAB* and *Gel* transcript expression were analysed in heterozygous *lms+/-;ap+/-* (**A-B’**) embryos, in single *lms-/-* mutant embryos (**C-D’**) and in double *lms-/-;ap-/-* mutants (**E-F’**) by *in situ* hybridisations with *dCryAB* or *Gel RNA* probes. (**A-B’**) Both *dCryAB* and *Gel* are prominently expressed in LT muscles of heterozygous *lms+/-;ap+/-* embryos. By contrast, *dCryAB* transcript levels are reduced in LTs (underlined area) in simple *lms-/-* mutants (**C-C’**) compared to (**A, A’**). Note that in *lms* mutants, *dCryAB* expression in DT1 remains unaffected (arrow). *dCryAB* down-regulation in a few remaining LTs is even more apparent (arrowheads) in double *lms-/-;ap-/-* mutants in which *dCryAB* expression in SBM and in DT1 is still detectable (arrows). *Gel* LT expression in the *lms-/-* context has almost completely vanished (underlined area) in **D** compared with (**B**), whereas its expression in the fat body (FB) remains high. No LT associated *Gel* expression could be detected in double *lms-/-; ap-/-* mutants. Note that gel expression in FB could still be detected (**F**). Conversely, whole muscle ectopic expression of *ap* using *Mef2-Gal4* driver is enough to induce *Gel* expression in all muscles (arrowhead points to LT muscles and arrows point to *Gel* ectopic expression in ventral muscles)(**G-H’**). **I**. Schematic representation of *dCryAB* and *Gel* transcriptional regulation.

### *dCryAB* promotes regular shapes and prevents branching of growing LT muscles

To test whether *dCryAB* helps set LT muscle features, we generated null allele (*dCryAB^HR^*) using CRISPR mutagenesis (Fig. 3A). *dCryAB* loss-of-function turned out to be homozygous lethal with mutants surviving until the late 3^rd^ larval instar. Compared to wildtype, the late stage embryos devoid of *dCryAB* (Fig. 3B-D) showed dissociations between LT1 and LT2 and/or LT2 and LT3 muscles (32% of segments) and irregular growth of LTs (28% of segments). The partial dissociation of LTs was in several instances associated with the irregular LT shapes (Fig. 3B,C) and branched LT muscle extremities (8% of segments) (Fig. 3C, right panel). We also noted a reduced number of LTs in 6% of segments (Fig. 3D). Branched LT muscles are also occasionally detected in *ap* and *lms* mutant contexts (Fig. S3) indicating that dCryAB is involved in preventing LTs branching downstream of LT iTFs.

**Figure 3.**
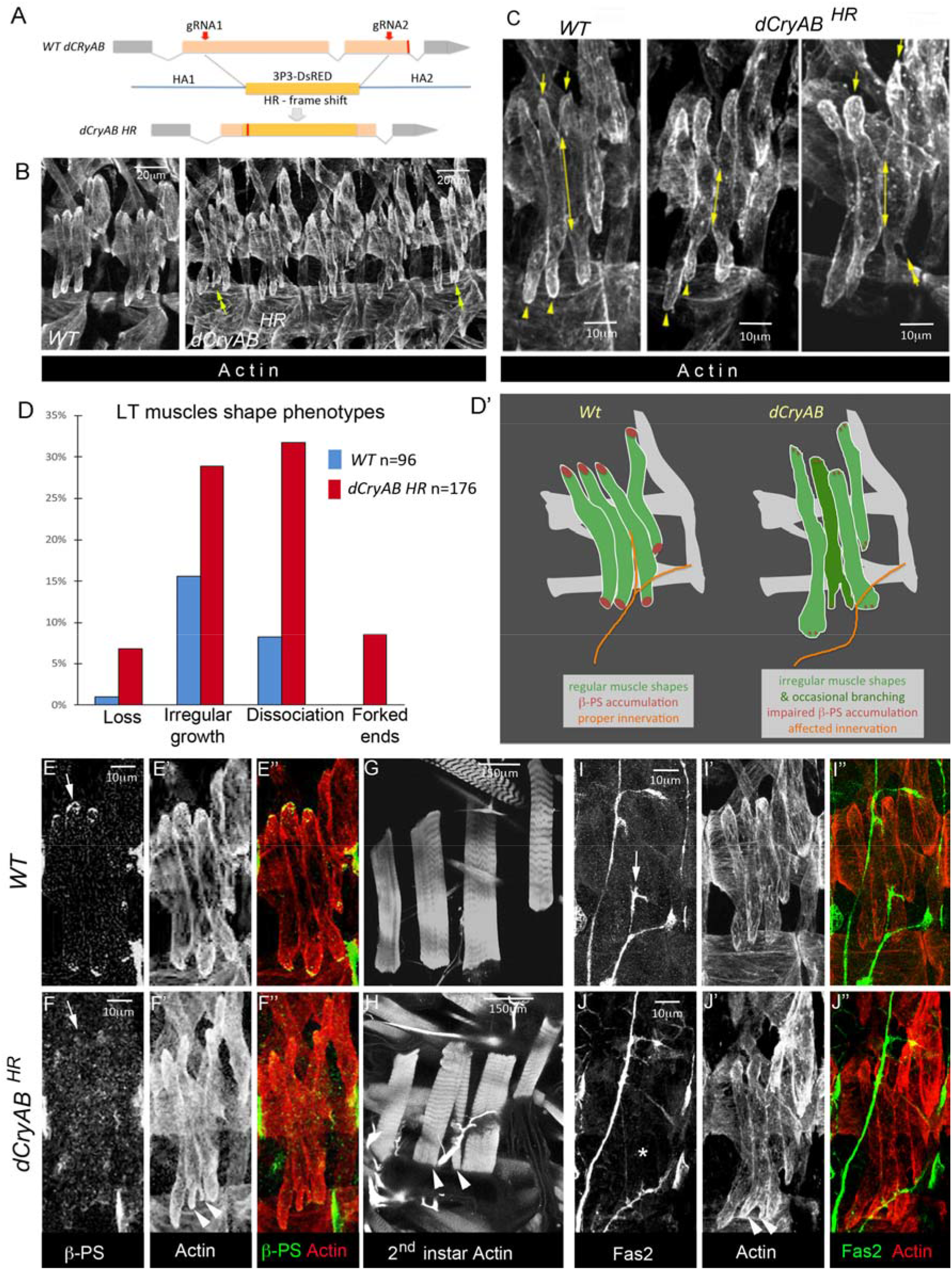
Aberrant growth-related LT muscle properties in *dCryAB* mutant embryos. **A.** Schematic representations of CRISPR mutagenesis of *dCryAB* locus. gRNAs targeting sites are indicated by red arrows, and positions of resulting stop codons by red lines. Homology directed recombination (HDR) with replacement of the *dCryAB* coding sequence by a 3P3-dsRed cassette was applied to generate the *dCryAB^HR^* null mutant line. **B-C**. Representative lateral views of muscles from late stage 15 embryos stained with anti-actin antibody to reveal muscle shapes. Yellow arrows and yellow arrowheads point to dorsal and ventral LT termini, respectively. They are aligned in wild type but appear at different levels in two neighboring LTs in *dCryAB^RH^*. suggesting irregular LT growth. Double-headed arrows show extents of contact between LT3 and LT4 muscles at the end of stage 15, which are often reduced in the *dCryAB^RH^* mutant context, indicating partial dissociation of growing LTs. Double-headed arrowhead points to forked/branched LT ends. **D.** Quantification of LT muscle phenotypes in *dCryAB^RH^* embryos presented as percentage of segments affected (*n* – number of abdominal segments examined). **D’.** A scheme representing LT muscle defects in *dCryAB* loss of function context. **E-F’’**. Lateral view of late stage 15 embryos, showing reduced □-PS Integrin staining at LT muscle termini in the *dCryAB^RH^* mutant context (**F-F”**) compared to WT embryo (**E-E”**). **G-H**. Lateral view of 2^nd^ instar *dCryAB^RH^* larvae, showing forked end phenotype on LT2 muscle (arrowheads in **H**). **I-J”**. Lateral view of late stage 15 embryos, stained for fasciclin 2 to show aberrant defasciculation of SNa motoneuron in *dCryAB^RH^* context (compare arrow (**I**) and star in **J**).

The irregular LT growth in *dCryAB* mutant embryos raised the question of whether *dCryAB* could impact on LT interactions with tendon cells and with motor neurons. By the end of embryonic stage 15, □-PS integrin accumulates at the extremities of muscles and at the surface of their cognate attachment sites, promoting the formation of myotendinous junctions (MTJs). Accordingly, in wild type stage 15 embryos, □-PS integrin labels ventral and dorsal ends of LTs (Fig. 3E-E”). However, in *dCryAB* mutants this □-PS accumulation was hardly detected, suggesting impaired MTJs formation (Fig. 3F-F”). Because in late *dCryAB* mutant embryos we do not detect an LT muscle detachment phenotype, observed in the □*-PS* loss-of-function context, we conclude that *dCryAB* mutation impairs but does not prevent □-PS accumulation at LT MTJs. Consistent with this, in 2^nd^ instar *dCryAB* mutant larvae, LTs, including those with branched ends, appear attached (Fig. 3G,H).

We then tested whether the LTs in *dCryAB* mutants were properly innervated. In wild type embryos, the dorsal branch of the SNa nerve innervating LTs defasciculates at the level of LT2 and grows dorsally within a gap between the ventral extremities of LT2 and LT3 (Fig. 3I-I”). In *dCryAB* mutants, most segments have wild type LT innervation, but in those with forked LT ends (essentially seen at the ventral extremity of LT2), the gap between LT2 and LT3 is filled, preventing defasciculation of SNa and LT innervation (Fig. 3J-J”).

Thus, by coordinating LT muscle growth, dCryAB ensures timely accumulation of □-PS integrin and optimal LTs attachment. The capacity of dCryAB to control LT shapes and prevent their branching facilitates proper LT innervation.

### *Gel* mutant embryos bear an increased number of LTs through fibre splitting

The CRISPR mutagenesis in the 5’ region of *Gel* resulted in two different null mutations, *Gel9.3* and *Gel9.8* (Fig. 4A), both leading to a premature stop codon. Because *Gel9.3* and *Gel9.8* exhibited similar molecular lesions, were homozygous viable, and showed equivalent LT muscle phenotypes (Fig. 4B), we chose one of them, *Gel9.3*, for further analyses.

**Figure 4.**
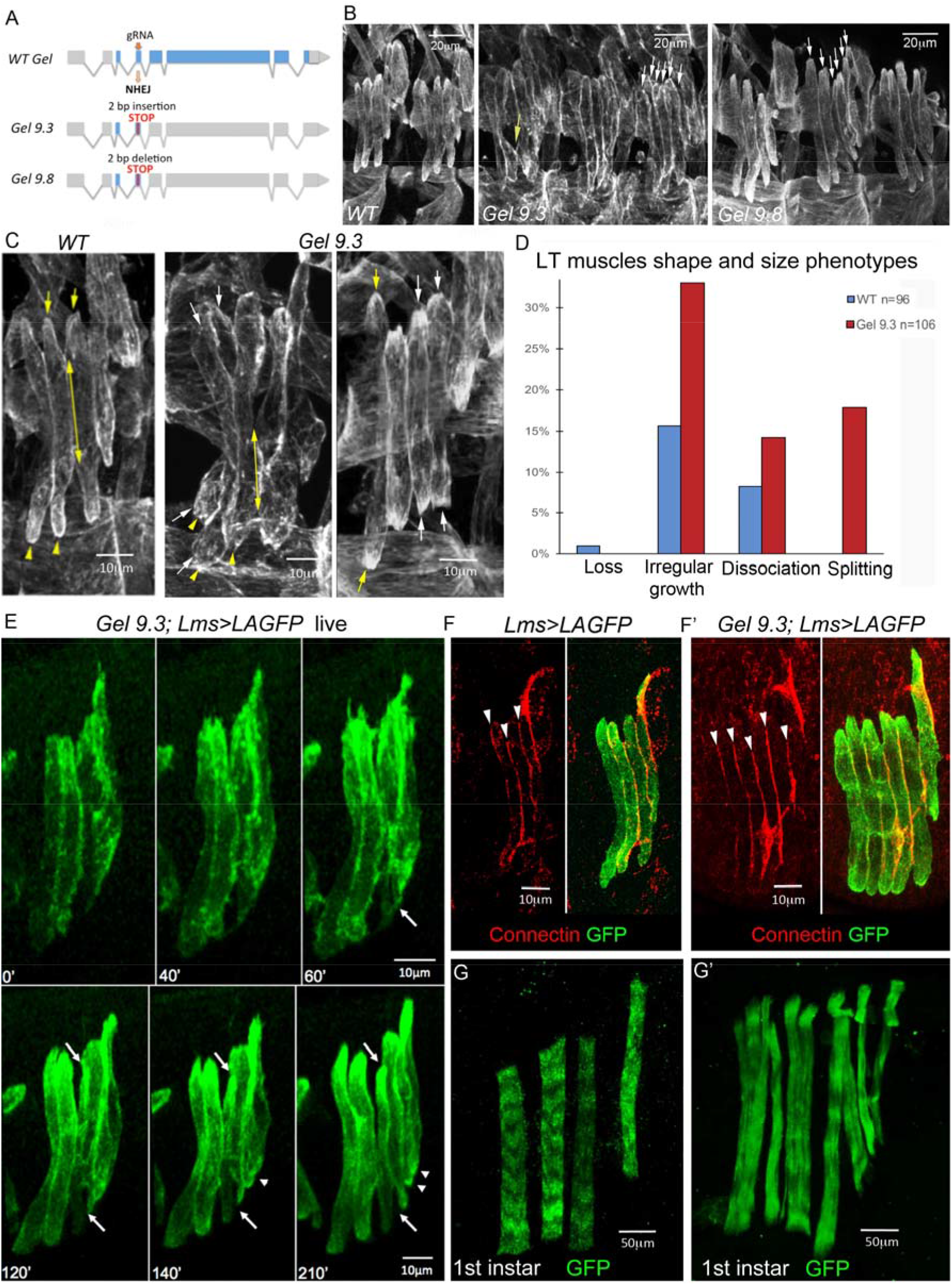
*Gel* mutant embryos show LT muscle splitting. **A**. Schematic representations of CRISPR mutagenesis of *gel* locus. The gRNA targeting site is indicated by red arrow and positions of resulting stop codons by red lines. *Gel 9.3* mutant was generated by a 2bp insertion and *Gel 9.8* mutant by 2bp deletion in the 5’ region of the *Gel* locus. **B-C**. Representative lateral views of muscles from late stage 15 embryos stained with anti-actin antibody. Upper panels (**B**) show two wild type abdominal hemisegments and three hemisegments from *Gel 9.3* and *Gel 9.8* mutants. White arrows point to excessive LT muscle extensions and the yellow arrow shows dissociation phenotype. Lower panels (**C**) are the zoomed views of LT muscles. Yellow arrows and yellow arrowheads point to dorsal and ventral LT termini, respectively. Double-headed arrows show extents of contact between LT2 and LT3 muscles at the end of stage 15, often reduced in the *Gel 9.3* mutants. White arrows point termini of muscles showing splitting phenotypes. **D.** Quantification of LT muscle phenotypes in *Gel 9.3* embryos presented as percentage of segments affected (*n* – number of abdominal segments examined). **E.** Snapshots over time of LT muscle shapes in *Gel 9.3* mutants revealed using the Lms>LifeActin-GFP LT sensor line. After 60 min, a first separation of muscle is observed (white arrow) on the ventral side of the LT3 muscle. This event is followed by actin cytoskeleton changes leading to progressive accumulation of actin on the dorsal side (upper white arrow) dissociation from LT2 muscle and individualisation of a new muscle fibre. Arrowheads show additional partial splitting of LT4 muscle on its ventral side. **F-F’.** Connectin-stained LT fibres in wild type and in *gel* mutants. Note increased number of connectin lines (arrowheads) in the segment with split LT muscles in the *gel* context (**F**) compared to the *Lms>LAGFP* context (**F**). **G-G’**. Lateral view of 1st instar larvae, showing split LT muscles with disorganised sarcomeric structures in the *Gel 9.3* context.

Despite irregularity in LT growth (32% of segments) and some LT dissociation phenotypes (14% of segments), *Gel* mutants also showed an increased number of LT muscles (17% of segments) (Fig. 4B-D), a phenotype previously described (6) and detected in particular in *ap* mutants (Fig. S3B), however not observed in the *dCryAB* loss of function context (Fig. 3).

To follow formation of supplementary LTs *in vivo*, we recombined *Gel9.3* with the *Lms>LifeActinGFP* (*LAGFP*) LT sensor line. We first confirmed that *Gel9.3;Lms>LAGFP* embryos form the supernumerary LTs, which become individualised with connectin-labeled cellular membranes (Fig. 4F,F’). We then performed time lapse experiments encompassing mid-embryogenesis and observed that supernumerary *Gel*-devoid LTs formed by fibre splitting. In the case presented (Fig. 4E, Fig. S4), the LT3 grows asynchronously, expands and subdivides progressively into two fibres with separate extremities. This aberrant morphogenesis appears to have a functional impact, as at the beginning of larval life, the striated sarcomeric pattern of split LT muscles is severely impaired (Fig. 4G,G’), indicating that their contractility is compromised. Thus, the supernumerary LTs in *Gel* mutants arise from enlarged fibres that eventually split. Another, interesting feature is that split LTs extremities accumulate □PS-integrin (Fig. 5A), suggesting that LT identity information ensuring choice of attachment sites is transmitted during splitting.

**Figure 5.**
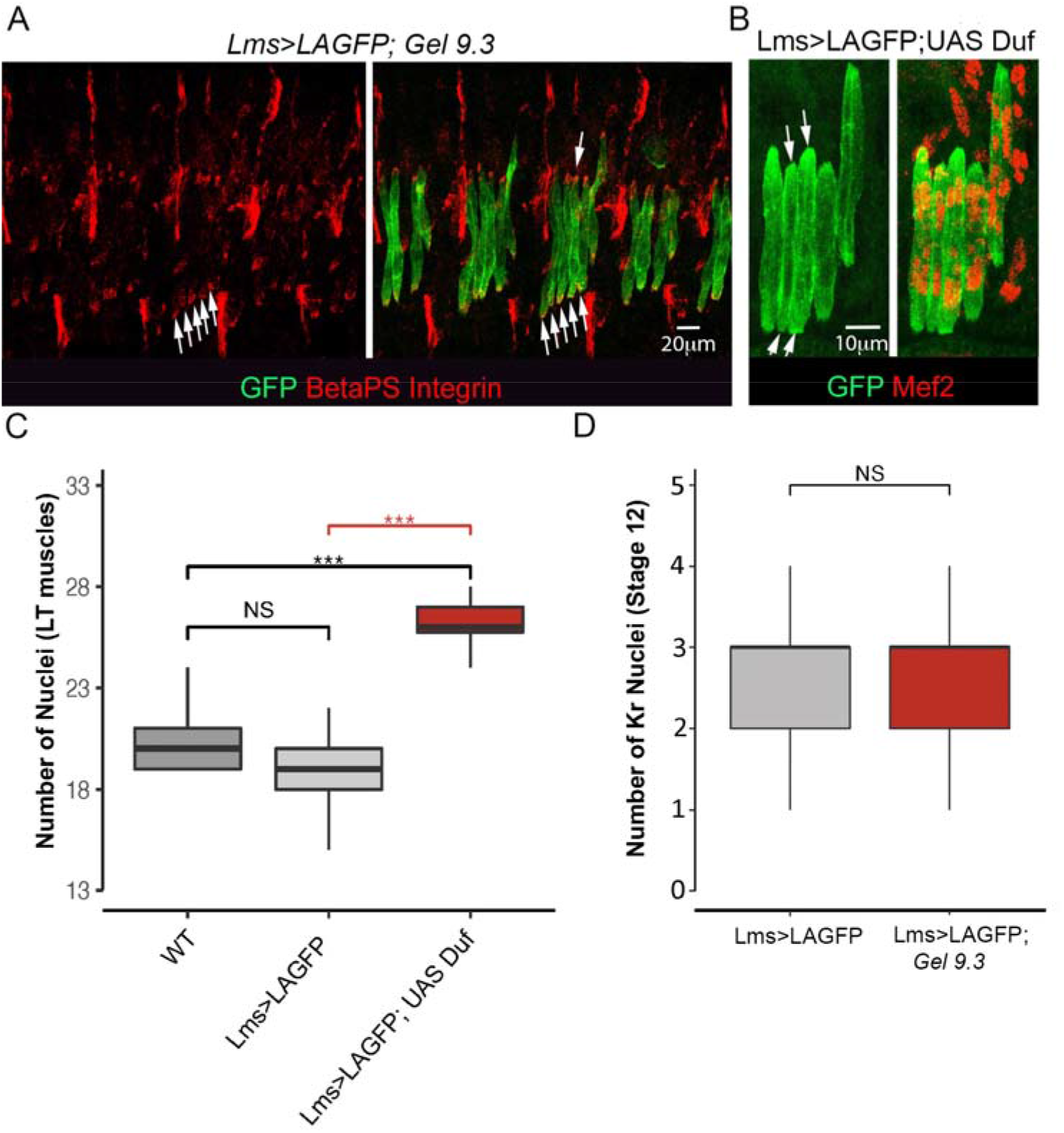
Split LT fibers keep LT properties but do not arise from an excess of progenitors and could be generated by local increase of myoblast fusion. **A.** Splitting in *Gel* mutants is associated with accumulation of □PS integrin (arrows) at termini of split LT fibres. **B-C.** *Duf* overexpression mimics the *Gel* LT splitting phenotype (arrows in **B**), which correlates with increased numbers of myonuclei. Myonuclei were counted (**C**) in 13 wild type abdominal segments and in 13 *Lms>LAGFP>Duf* abdominal segments with splitting events and in a larger number of abdominal segments (*n* = 43) from *Lms>LAGFP*. A Wilcoxon-Mann-Whitney test was applied to assess statistical significance. \*\*\**p* < 0.001. **D.** Graphical representation of number of Kr progenitors at stage 12 showing that supernumerary LT muscle fibres are not due to an excess of LTs progenitors. A large number of hemisegments in *Lms>LAGFP (n* = 150) and *Gel 9.3;Lms>LAGFP* mutant embryos (*n* = 114) were analysed. A Wilcoxon-Mann-Whitney test was applied to assess statistical significance. \*\*\**p* < 0.001

### *Gel* controls LT muscle size by preventing excessive myoblast fusion

On performing time lapse experiments, we observed that splitting occurred during the developmental period in which muscle fibres grow by fusing with surrounding myoblasts, and that split LTs were enlarged compared to non-split neighbours (Fig. 4E). Our previous findings showing that muscle size depends on the number of fusion events [15, 24] thus raised the question of whether LT splitting could be associated with increased fusion. This appears to be the case since the LT-targeted increase in fusion by overexpressing *Duf* could lead to splitting (Fig. 5B,C). On the other hand we found that *Gel* is expressed in developing LT muscles but not in FCs (Fig. S2) and that the number of LT FCs in *Gel* mutants remains unchanged (Fig. 5D), suggesting that the supernumerary *Gel*-devoid LTs arise by an aberrant fusion-involving muscle morphogenesis.

Indeed, the number of Mef2-positive nuclei in LT1-LT4 at embryonic stage 16 was significantly higher in *Gel* mutants including the transheterozygous *Gel9.3/Gel9.8* context compared to controls (Fig. 6A-D). We then sought to determine whether the LT-specific increase in fusion events observed in *Gel* mutants could impact on the fusion programs of neighbouring muscles. The number of myonuclei in immediate LT neighbours in the SBM and LO1 muscles (but not in more ventrally located VT1) was reduced, indicating that local availability of FCMs could impact on fusion programs (Fig. 6E,F). Muscle splitting observed in *Gel* mutants thus results from excessive fusion, with late fusion events that could be detected associated with LTs showing a split phenotype (Fig. 6B, right panel). The capacity of *Gel* to negatively regulate fusion was confirmed by the reduced number of myonuclei in LTs in which *Gel* was prematurely activated (Fig. 6D) and by the large number of unfused myoblasts seen in embryos with ectopic *Gel* expression in all muscles (Fig. 6G). Thus, we propose that *Gel* triggers fusion arrest in LTs. In *Gel* loss of function context LTs continue to grow by fusion and eventually split (Fig. 6H).

**Figure 6.**
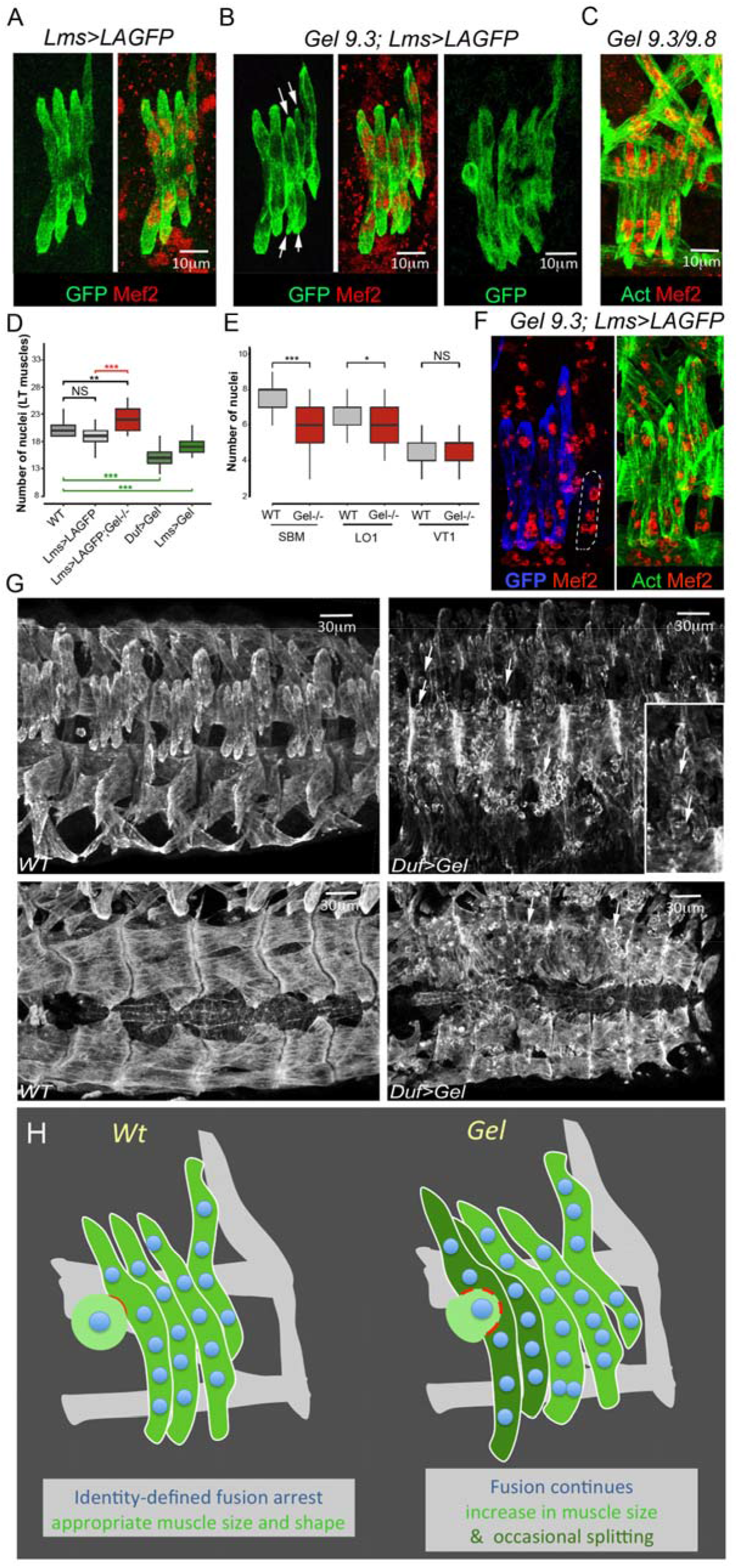
LT muscle splitting in *Gel* mutant embryos is due to an excessive myoblast fusion. LT muscle shapes at late stage 15 revealed by anti-GFP staining in Lms>LifeActin-GFP sensor line (**A**), in *Gel 9.3* context (**B**) and in transheterozygous *Gel 9.3/9.8* embryos (**C**). Myonuclei are visualised using anti-Mef2. Arrows indicate LT splitting in *Gel* homozygous (**B**) and the transheterozygous context (**C**). Late fusion events are observed in *Gel 9.3;Lms>LAGFP* embryos (**B**, right panel). (**D**) Number of Mef2 positive LT1-LT4 nuclei is increased in *Gel* mutants (*n* = 30) compared to wild type (*n* = 13) and *Lms>LAGFP* contexts (*n* = 43). An opposite effect (reduced number of nuclei) is observed in LTs of Duf>Gel (*n* = 80) or Lms>Gel embryos (*n* = 65) (green boxplots). A Wilcoxon-Mann-Whitney test was applied to assess statistical significance. \*\*\**p* < 0.001 (**E**). Excessive fusion in *Gel* mutant with split LTs induce a decrease in the number of nuclei in neighbouring SBM and LO1 but not in ventrally located VT1 muscle. For SBM: WT (*n* = 66), *Gel 9.3 (n* = 110), for LO1: WT (*n* = 58), *Gel 9.3 (n* = 100), for VT1: WT (*n* = 68), *Gel 9.3 (n* = 93). A Wilcoxon-Mann-Whitney test was applied to assess statistical significance. \*\*\**p* < 0.001. (**F**) Reduced number of nuclei in SBM muscle (dotted line) of *Gel 9*.3 embryo (four instead of seven in the *wild type*). **G.** Lateral and dorsal views of anti-actin stained stage 16 *wild type* and *Duf>Gel* embryos (right panels) showing myoblast fusion defects. Arrows point to unfused myoblasts. **H.** A scheme of *Gel* loss of function phenotypes.

## Discussion

TRAP was first developed to isolate polysome-associated mRNA from a subset of neurons in mice [16] and was later adapted to other model organisms including *Xenopus* [25], zebrafish [26] and *Drosophila* [17, 27]. Here we applied TRAP to determine the first translatomic signatures underlying diversification of muscle types, and we identified *Gel* and *dCryAB* as new LT muscle identity realisator genes. *dCryAB* contribute to preventing LTs branching, and *Gel* plays a role in LT muscle size control by limiting the number of fusion events. Consequently, supernumerary myonuclei are present in *Gel*-devoid LTs, which eventually split. Both splitting and branching are low penetrance phenotypes (17% and 8% of segments, respectively), indicating that *Gel* and *dCryAB* are not the sole identity realisators that prevent these aberrant growth-related LTs features.

Formation of branched muscle fibres has recently been reported as a result of adversely affected muscle identity [28], and supernumerary LTs were also detected in *ap* and *lms* mutant embryos [6], indicating that a muscle identity-dependent shape and size control system must exist in developing muscles.

In humans, *CryAB* mutations are associated with desminopathies in which aberrant muscle fibres with branched morphology are frequently detected [29]. *Gelsolin* mutations cause amyloidosis, characterised by the toxic accumulation of protein aggregates, which can lead to an inclusion body myositis (IBM)-like phenotype with necrotic and centronuclear split fibres [30, 31].

Functional analyses of *dCryAB* and *Gel* in *Drosophila* embryos indicate that muscle fibres branch or split when the identity realisators for these muscle are not properly activated. Reduced levels of □PS-integrin at the extremities of *dCryAB*-devoid LTs suggests weak interactions with attachment sites and could explain observed LTs overgrowth and dual attachment of branched fibers. These *dCryAB* loss of function phenotypes could result from inappropriate actin cytoskeleton dynamics in growing LTs and/or affected function of cheerio/filamin, a direct dCryAB interactor [23].

In advanced stages of muscular dystrophies, a large subset of muscle fibers shows longitudinal splitting, described already more than forty years ago (34). After chemical or physical injury, split fibers form also in undergoing regeneration *wild type* muscles (35) suggesting a link between myoblast fusion-involving regeneration (chronic in dystrophic context) and muscle splitting. However, despite critical impact on dystrophic muscle cytoarchitecture, on vulnerability to contraction-induced damage and on muscle weakening, mechanisms of muscle fibers splitting remain unexplored (36; 37).

Our finding in *Drosophila* that local increase in fusion occuring in *Gel* mutant embryos causes split fiber phenotype points to the excessive fusion as a prerequisit of splitting.

## Acknowledgements

This work was supported by AFM-Téléthon (MyoNeurAlp Strategic Program), Agence Nationale de la Recherche (Tefor Infrastructure Grant), ANR JC (Cardiac-SPE) and Fondation pour la Recherche Médicale (Equipe FRM Award). We thank the staff of Genecore EMBL Heidelberg for their technical assistance.

## Materials and methods

### Fly stocks

All *D. melanogaster* stocks and crosses were grown on standard medium at 25°C. The following strains were used: *ap^UGO35^* (gift of J. Botas, Baylor, Houston, USA), *lms^s95^* (gift of D. Müller, Universität Erlangen-Nürnberg, Germany), *UAS-LifeAct-GFP* (Bloomington, 35544) and *UAS-Dumbfounded* (*UAS-Duf*) (gift of M. Ruiz-Gomez, Spanish National Research Council, Spain). To generate the TRAP lines, the *UAS-RpL10a-EGFP* line was crossed with the *Slou-Gal4* (gift of M. Frasch, Universität Erlangen-Nürnberg, Germany), *Lms-Gal4* (Janelia Farm collection, 46861) and *Duf-Gal4* (gift of K. Vijayraghavan, TIFR, India) driver lines to specifically target polysomes in the respective muscle populations.

### Generation of *gel and dCryAB* knock-out lines by CRISPR-Cas9

The different gRNAs were designed using CRISPR optimal target finder [32]. To generate *gel* guide we used the pair of primers 5’-GTCGAGACCTCGACCGATGAGGC-3’ and 5’-AAACGCCTCATCGGTCGAGGTCTC-3’. The primers were annealed, digested by Bbs1 and cloned into pCFD3 plasmid (plasmid #49410, Addgene) for *gel*. For *dCryAB* a HDR-based CRISPR technology was applied. First, two guides targeting 5’-CTTGGACCAGCACTTCGGTC-3’ and 5’-GGAGGACAACGCCAAGAAGG-3’ sequences located in the 5’ and 3’ regions of the *dCryAB* gene, respectively, were designed and cloned by Gibson assembly into pCFD4 plasmid (#49411). The homology arms HA1 of 1045bp and HA2 of 1058bp were amplified using the following pairs of primers:

Forward HA1: 5’-ATATCACCTGCATATTCGCAGCGACGTCATCTCTTTCGTCTG-3’
Reverse HA1: 5’-ATATCACCTGCATATCTACAAGAGGCGCGAGGTGCGCATTG-3’
Forward HA2 : 5’-ATATGCTCTTCATATAGGTGGAGACCTCCACCGCC-3’
Reverse HA2 : 5’-ATATGCTCTTCAGACTTCGTCAGGTTCGGTTACTCCG-3’ into AarI and SapI cloning sites of donor pHD-DsRed plasmid (#51434). Plasmids were injected into Nos-Cas9 embryos (BestGene).

To establish KO lines, molecular characterisation of target loci was performed as described [33]. Briefly, genomic DNA was extracted from individual larvae by crushing them in 20 μL QuickExtract solution (Cambio) and releasing the DNA in a thermomixer according to the supplier’s instructions. We used 1 μL (previously diluted five times) of the supernatant in 25 μL PCR reactions. PCR products were then sequenced by Sanger. Indels can be observed as regions with double peaks in heterozygous flies, corresponding to wild type and mutated allele respectively. In the case of *dCryAB*, homologous recombination events were recovered by selecting flies with red eye fluorescence.

### RNA extraction and RT-qPCR

mRNA was extracted from Lms, Slou and Duf muscle populations using TRIzol reagent (Invitrogen) following the manufacturer’s instructions. RNA quality and quantity were then assessed using Agilent RNA 6000 Pico kit on Agilent 2100 Bioanalyzer (Agilent Technologies).

### Microarray analysis

Agilent 8×60K probe (60-mer) gene expression microarrays were used. We assessed the quality of triplicates using Pearson’s correlation test. Correlations for all conditions were 85% or higher. The microarray data were quantile-quantile normalised. Gene expression data from Slou-, Lms- and Duf-positive cells were compared to the whole embryo datasets to generate lists of genes differentially expressed, fold change ≥ 2, *p* < 0.05. GO Princeton software was used to assign GO classification. We then compared Lms- and Slou-positive cells to Duf-expressing cells to make two lists of differentially regulated muscle-specific genes, fold change ≥ 2, *p* < 0.05. We computed and compared GO biological processes from these two lists using an R package cluster profiler.

#### Temporal transition heatmap

Translational temporal profiles from Slou and Lms at the three time points were converted to “transition values” defined as log ratios between T2 and T1, T3 and T2. Transition values were considered as three discrete classes: upregulated (>1), stable (between −1 and 1), and downregulated (<1). Thus expression profiles from Lms and Slou muscles, which contain three temporal windows, were converted into vectors of two transitions (Tr1, Tr2), which allow the determination of correct gene behaviour. For example, the profile “red-red” group genes whose RNA level increases between T1 and T2 (Tr1), and then continues to increase between T2 and T3 (Tr2).

### *In situ* hybridisation and immunostaining

Embryos were dechorionated and fixed in 4% paraformaldehyde/heptane for all immunohistochemistry. Fluorescent *in situ* hybridisation with a TSA amplification system (Perkin-Elmer) and immunohistochemistry was as described previously [13]. To generate the RNA probe for *gel* (primers used: 5’-5’AATCGACTCCGTGGTGACTC-3’ and 5’-GGGAGGCCAAAGATGAGCTGTC-3’) the corresponding DNA sequences were cloned by PCR in pCR II topo vector. The corresponding anti-sense RNAs were transcribed *in vitro* using T7 or SP6 RNA polymerase. For *dCryAB*, Gold collection clone *GH01960* was used to generate RNA probes. For fluorescent staining, the following antibodies were used: rabbit anti-β3 tubulin (1:5000; R. Renkawitz-Pohl, Philipps University, Marburg, Germany), rat antiactin (1:300, MAC 237; Babraham Bioscience Technologies), rabbit anti-Mef2 (Nguyen HT, 1:2000) anti-GFP (1:1000 Developmental Studies Hybridoma Bank (DSHB)), mouse anti-Integrin βPS (1:200 DSHB) and mouse anti-connectin (1:200 DSHB). Cy3, Cy5, and 488 conjugated secondary antibodies were used (1:300; Jackson Immuno-Research). Embryos were mounted in anti-fade Fluoromount-G reagent (Southern Biotech). Labelled embryos were analysed using an Leica SP8 confocal microscope equipped with a HyD detector and a 40X objective. Images were processed with ImageJ.

### TRAP experiment

RPL10aGFP-tagged embryos were collected, and messenger RNAs from the different muscle populations isolated as described [17]. Micro-array data generated in this study were deposited to GEO database : GSE137443: GSM4079386 to GSM4079439

### Live imaging

The *Lms-Gal4; UAS-lifeAct* double transgenic line was generated and used for time lapse imaging of LT muscle formation in the *gel* mutant context. Image acquisition was performed on manually aligned living embryos at 21 °C using an inverted Leica SP8 confocal microscope. The time interval between acquisions was set to 3 min and the acquisition time was 3–4 h. Movies were generated and analysed using Imaris software (Bitplane).

