## Supplemental information for "*Gelsolin* and *dCryAB* act downstream of muscle identity genes and contribute to preventing muscle splitting and branching in *Drosophila*"

### Supplementary Figure legends:

**Figure S1. Sensitivity assessment of translational profiling by TRAP.**

**A-B.** Vulcano plots showing distribution of up- and down-regulated genes in Slou- and Lms-positive muscle subsets at the T3 time point. Note that genes known to be specifically expressed in the slou population (*slou, org-1*) and the lms population (lms) are consistently up-regulated. **C-D**. Distribution of Slou and Lms up-regulated and down-regulated genes from the T2 time window with respect to modEncode RNA expression level based on whole embryo RNA sequencing (stage 12–16).

**Figure S2. *Gel* is expressed in LTs from stage 14 and is present in other mesodermal tissus.** *Gel* expression in stage 13 (**A,A'**) and stage 14 (**B,B'**) embryos revealed by *in situ* hybridisation with the *Gel* RNA probe. Muscles are stained with anti-actin antibodies. *Gel* transcripts are first detected in LT muscles at stage 14 and are also found (**C,C'**) in visceral muscle (VM) and fat body (FB) (arrows).

**Figure S3. Branched and split phenotypes in LT iTF mutant embryos. (A-C)** Lateral views of four hemisegments of stage 16 embryos stained with anti-Actin (A, B) and anti-β3-Tubulin antibodies. (A) wild type muscle pattern. (B) An example of split (arrows) LT muscles in apterous $ap^{UGO35}$ mutant embryos. Asterisks point to lacking LT muscles (C) An example of branched LT muscle phenotype (arrowheads) in $lms^{S95}$ mutant embryo.

**Figure S4. *In vivo* visualisation of LT splitting in *Gel* mutants.**

Time lapse movie (3 h 30 min, with 3 min frame interval) from *Gel;lms>LAGFP* embryos showing one abdominal segment (A5) with developing LT muscles. The film begins at late stage 14 and ends at late stage 16 when first muscle contractions appear. Splitting of LT3 and LT4 muscles could be observed in segment A5. It is initiated at stage 15 and fully apparent with split fibre morphology and duplicated ends at stage 16.

Figure S1

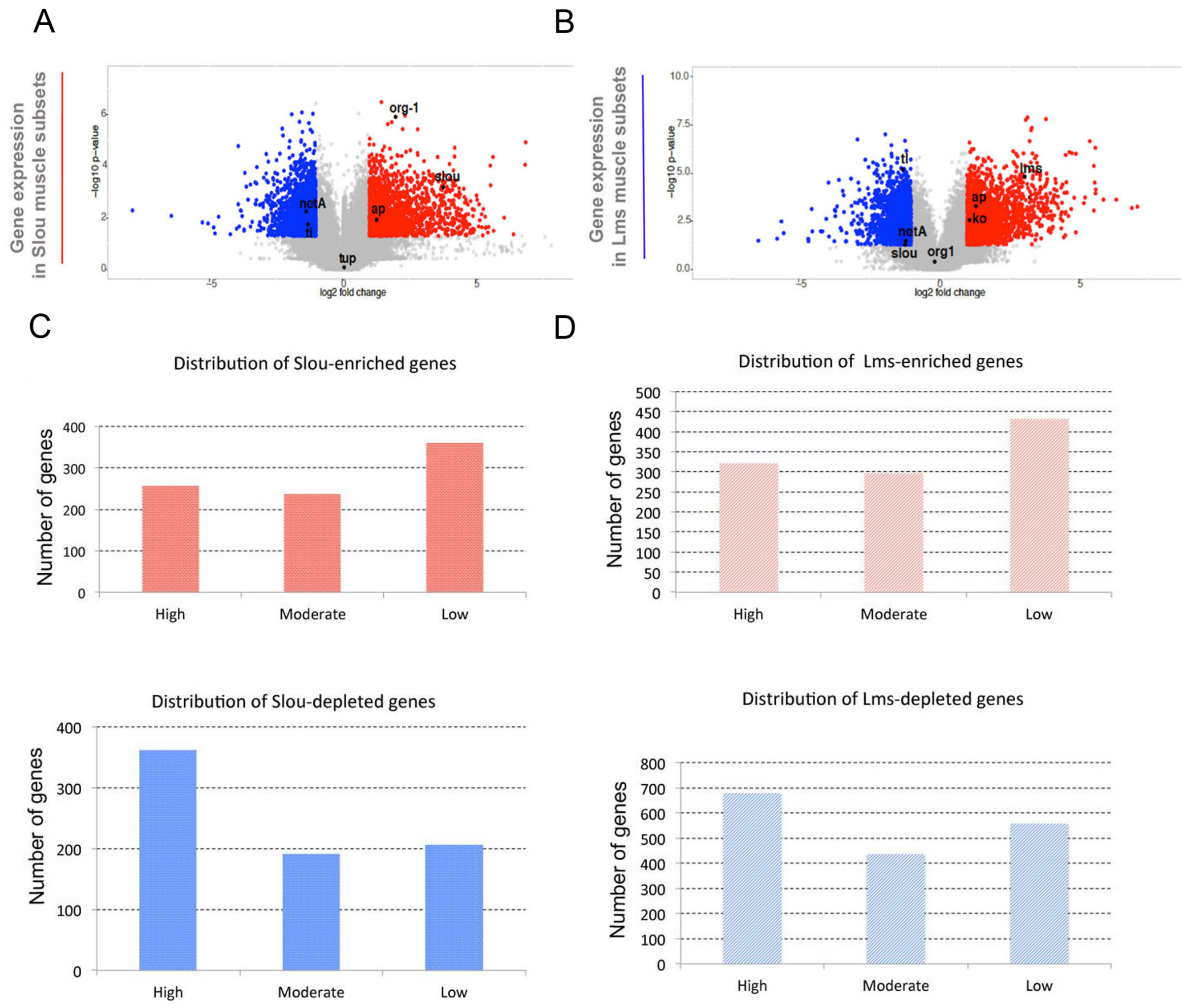
A
B
Gene expression in Slou muscle subsets
-log10 p-value
log2 fold change
org-1
slou
netA
ap
tl
tup
Gene expression in Lms muscle subsets
tl
lms
ap
ko
netA
slou
org1
C
D
Distribution of Slou-enriched genes
Number of genes
High
Moderate
Low
Distribution of Lms-enriched genes
Distribution of Slou-depleted genes
Distribution of Lms-depleted genes

Figure S2.

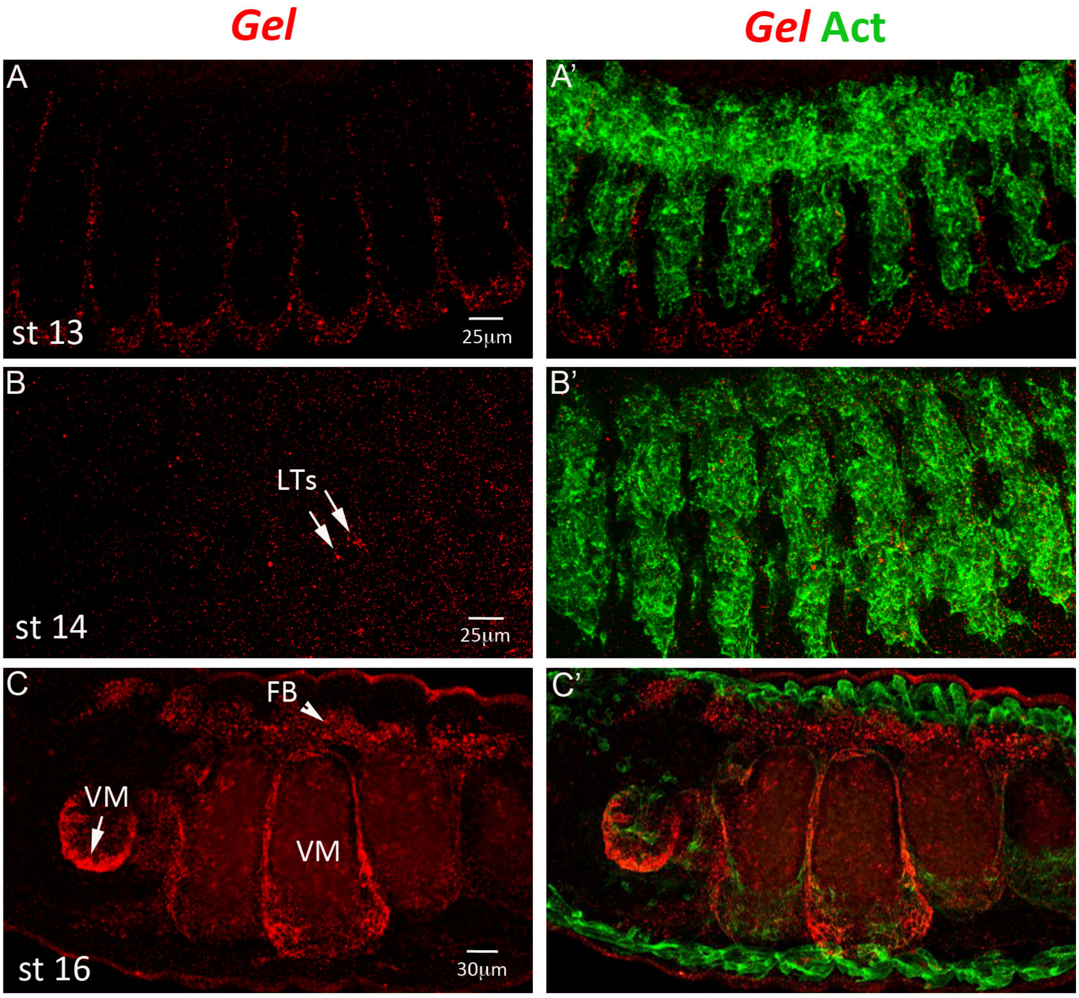

Figure S3

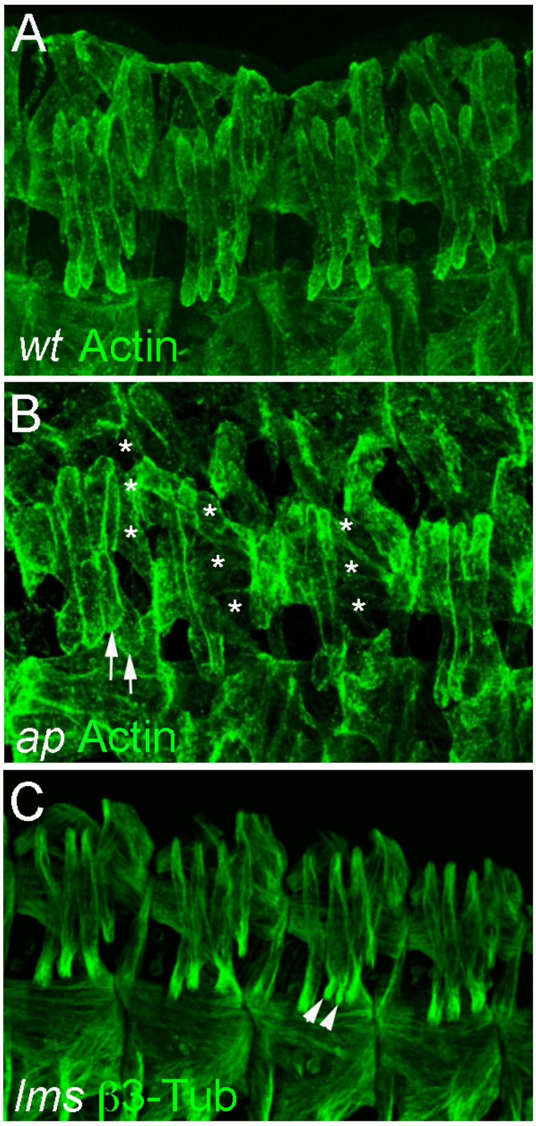
